# Stratified Palliobulbar and Midbrain Circuits Control the Timing and Acoustics of Learned Vocalizations

**DOI:** 10.64898/2026.10.26.761713

**Authors:** Ziran Zhao, Todd F. Roberts

## Abstract

In mammals and birds, production of learned vocalizations depends on prominent projections from the pallium to the brainstem^1–8^. Although pallial and cortical coding mechanisms for speech and birdsong are becoming increasingly clear^9–19^, understanding how palliobulbar projections supervise brainstem circuits has remained challenging. Here we identify the functional organization of pallial and brainstem control of learned song in the zebra finch, a songbird in which genetic control of neuronal pathways is now feasible^18,20,21^. We find that palliobulbar projections to the medulla and midbrain periaqueductal gray (PAG) arise from largely non-overlapping neuronal populations. Genetic silencing of these pathways reveals that projections to the medulla control the learned acoustic content defining individual song syllables, while projections to the PAG control the respiratory timing needed to sequence syllables. Using large-scale neuronal recordings, we find that song and innate calls are uniquely encoded within the PAG. Dual recordings from the palliobulbar neurons and the PAG suggest hierarchical encoding of expiratory and inspiratory events along this pathway. Lastly, genetic silencing reveals that PAG projections to the medulla are needed for birds to generate learned song and innate calls. Thus, stratified palliobulbar pathways control respiratory-timing and acoustics for song. Yet, without the PAG, they are insufficient to produce any vocalizations.

## Main

Vocalizations are used by many vertebrates to communicate internal states, facilitate social exchanges, coordinate reproductive behaviors and mediate parent-offspring interactions ^22–24^. Among vocalizing species, a small and phylogenetically diverse group of animals evolved the ability to culturally transmit their vocalizations across generations through imitation (**Fig. 1a**), a skill foundational to speech and language, as well as birdsong, parrot contact calls, and the songs and calls of whales and seals^25^. The production of learned vocalizations requires extreme dexterity and is associated with hypertrophied pallial circuits, which are thought to have extended direct connections with premotor respiratory neurons and vocal organ motor neurons in the medulla^1–5,26–28^ (collectively referred to here as vocal-motoneurons (V-MNs); **Fig. 1b-c**). Although studies across humans, parrots, and songbirds have shed light on how learned vocalizations are represented in the cortex and avian pallium^9–19^, the functional logic for how palliobulbar and brainstem pathways execute vocal-motor commands is not known.

**Fig. 1.**
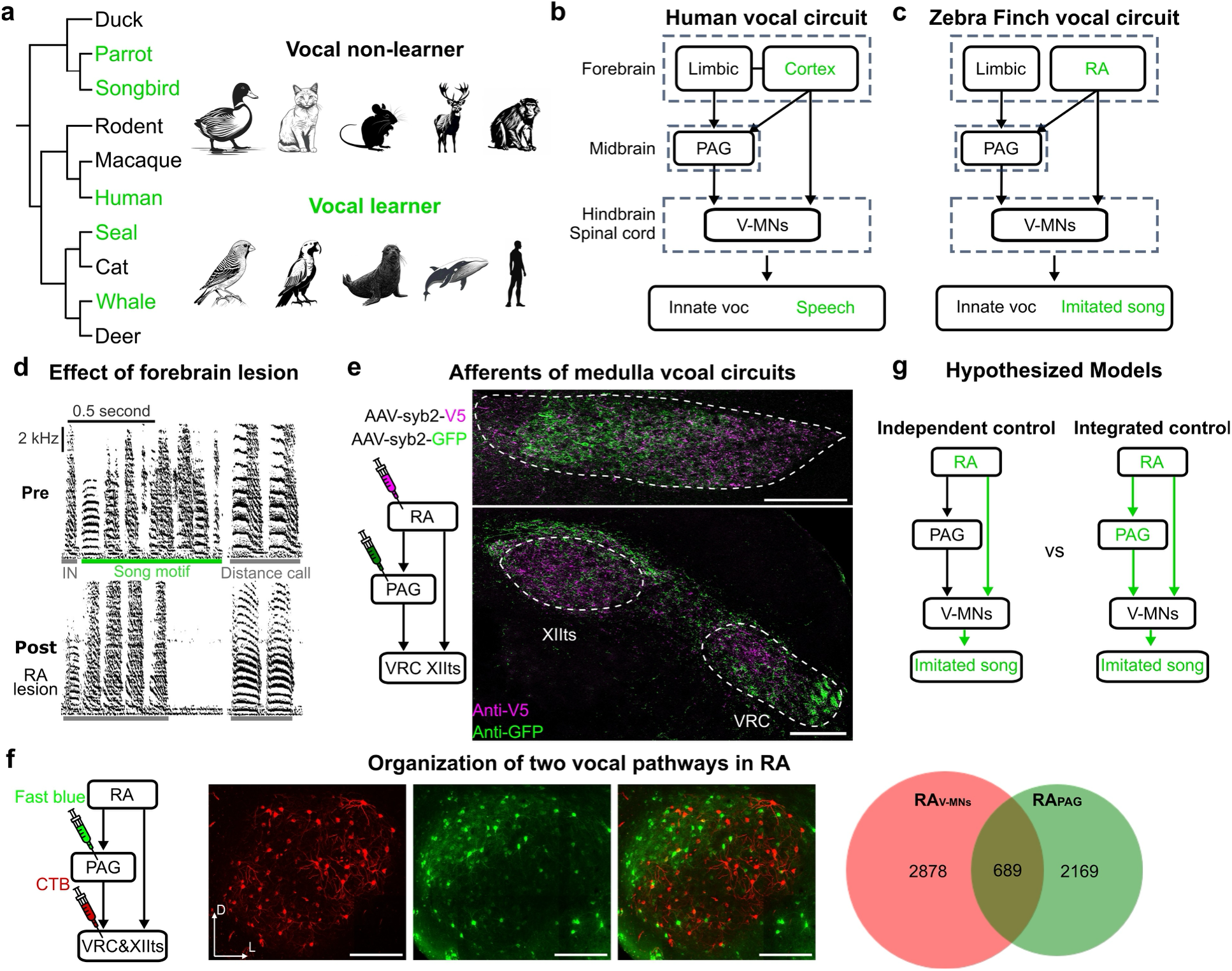
Descending neural circuits regulate innate and learned vocalizations. **(a)** Vocal non-learners and vocal learners across the phylogenetic tree. Skilled vocal learning species are highlighted in green. The cartoon illustration of animals was created using PIXLR (https://pixlr.com). **(b and c)**, Schematic of human (b) and zebra finch (c) descending vocal circuits. The brain regions known to be necessary for learned vocalization are marked in green. **(d)** Song spectrograms of innate and learned vocalization before and after RA lesion (top and bottom). The song motif is underlined in green, and innate vocalizations, introductory notes (IN) and distance calls, are underlined in grey. n=3 animals. **(e)** Viral tracing of the hindbrain vocal circuit. Left, the schematic of the viral injection. Right, anti-V5 is labelled in magenta and anti-GFP is labelled in green. Green and magenta fluorescence in the hindbrain indicate the PAG and RA presynaptic sites, respectively. The dashed contours outline PAG, VRC and XIIts. n=3 animals. Scale bars, 200 μm. **(f)** Two RA populations exclusively projecting to PAG and the hindbrain. Left, circuit schematic of tracer injection. Medial, cells labelled in red or green are RA neurons projecting to the hindbrain or PAG, respectively. Right, Venn diagram of V-MNs/PAG-projecting RA neurons are plotted with data collected from three animals. **(g)** Circuit schematic of two hypothesized models. Song-related brain regions are in green. Arrowheads or flat bars indicate the overall effect of the projection on the downstream target is excitatory or inhibitory, respectively.

Songbirds provide an important model system for studying the neural circuits for learning and producing vocalizations^1^. Male zebra finches produce a learned song motif that is characterized by the serial and stereotyped production of several imitated song syllables. Their vocal repertoire also includes introductory notes that precede production of their song motifs and distance calls, both of which are innate but contain developmentally learned acoustic features^29–31^. The zebra finch motor pallium contains a specialized vocal-motor region, which is not seen in non-vocal learning birds, named the robust nucleus of the arcopallium (RA). Historically, research has shown RA to be necessary for production of learned song(s) and the learned features of introductory notes and distance calls^32^. RA is thought to control learned vocalizations through its topographically organized projections to V-MNs. RA also makes a prominent projection to the lateral periaqueductal gray (PAG)^33–35^, a highly conserved midbrain region that gates production of innate vocalizations in other species^23,36–42^. The lack of genetic methods in songbirds for pathway-selective manipulations has challenged the ability to dissect the function of this palliobulbar pathway. Consequently, the specific roles of RA projections to V-MNs and the PAG are not known, nor is the role of the PAG in production of learned vocalizations.

Here we combine pathway selective genetic silencing, large-scale Neuropixels recordings in freely singing birds, and functional circuit mapping to systematically examine the role of palliobulbar pathways and the PAG in the control of zebra finch song. We show that palliobulbar pathways have target-specific stratified control of the fine acoustic features and song syllable timing. Furthermore, we reveal that the midbrain PAG, not the palliobulbar pathway, is the final arbiter of whether birds can produce any vocalizations, including their learned song.

## RESULTS

As a starting point, we re-examined the basic function of the zebra finch vocal-motor pallium by making excitotoxic lesions of RA^32^ (n = 3 birds). This blocked the ability of birds to spontaneously produce their song. When presented with a female, male birds would exhibit courtship behaviors like approaches and beak wipes but mostly failed to produce any song syllables^32,43^. In successful attempts at singing (**Fig. 1d**), we observed highly disrupted behavior that lacked the learned timing (i.e. temporally organized onset and offset of song syllables) and modulation of spectral features (frequency modulation, harmonic content etc.) characteristic of the birds’ pre-lesion song. Likewise, although birds still produced introductory notes and distance calls, RA lesions eliminated learned acoustic features in these vocalizations. Thus, RA is critical in the production of song syllables and controls the spectral features of introductory notes and distance calls

### Two palliobulbar pathways control learned song

We next mapped the synaptic connectivity of palliobulbar projections from RA. We made viral tracing tools using a synaptobrevin-2 (Syb2) fusion protein to tag presynaptic release sites (**Fig. 1e**). Injections into RA revealed a strong density of synaptic release sites across V-MNs, including overlap with motoneurons in the tracheosyringeal division of the hypoglossal cranial nerve (nXIIts) and the more laterally positioned ventral respiratory column (VRC). In the midbrain we found a strong density of synaptic release sites in the lateral PAG, a portion of the PAG historically referred to as DM (dorsomedial nucleus of the intercollicular complex)^44,45^, which also innervates V-MNs in the medulla.

In humans, corticobulbar projections to medullary V-MNs and the vocal PAG are thought to arise from the laryngeal motor cortex and anterior cingulate cortex, respectively^3,4,46–49^. Therefore, we asked if pallial projections onto V-MNs and the PAG also arise from non-overlapping palliobulbar pathways in the songbird (**Fig. 1b,c and Extended Data Fig. 1a,b**). Injections of anatomical retrograde tracers in the PAG and V-MNs revealed that 88% of RA projection neurons exclusively project to either the PAG (RA_PAG_) or to medullary V-MNs, including the VRC (RA_V-MNs_; **Fig. 1f**). Thus, like us, the songbird has two largely separate palliobulbar projection pathways that may be used to control its learned vocalizations (**Fig. 1g**).

### Stratified palliobulbar control of song

The anatomical segregation of palliobulbar projections to the PAG and medulla presented the possibility to examine their individual roles in song production. We silenced each pathway using viral retrograde expression of Cre from either the PAG or medulla, and expression of Cre-dependent tetanus neurotoxin (TeNT) in RA^18,50–52^, which prevents presynaptic neurotransmitter release by blocking synaptic vesicle fusion (**Fig. 2a,b, and Extended Data Fig. 2**).

**Fig. 2.**
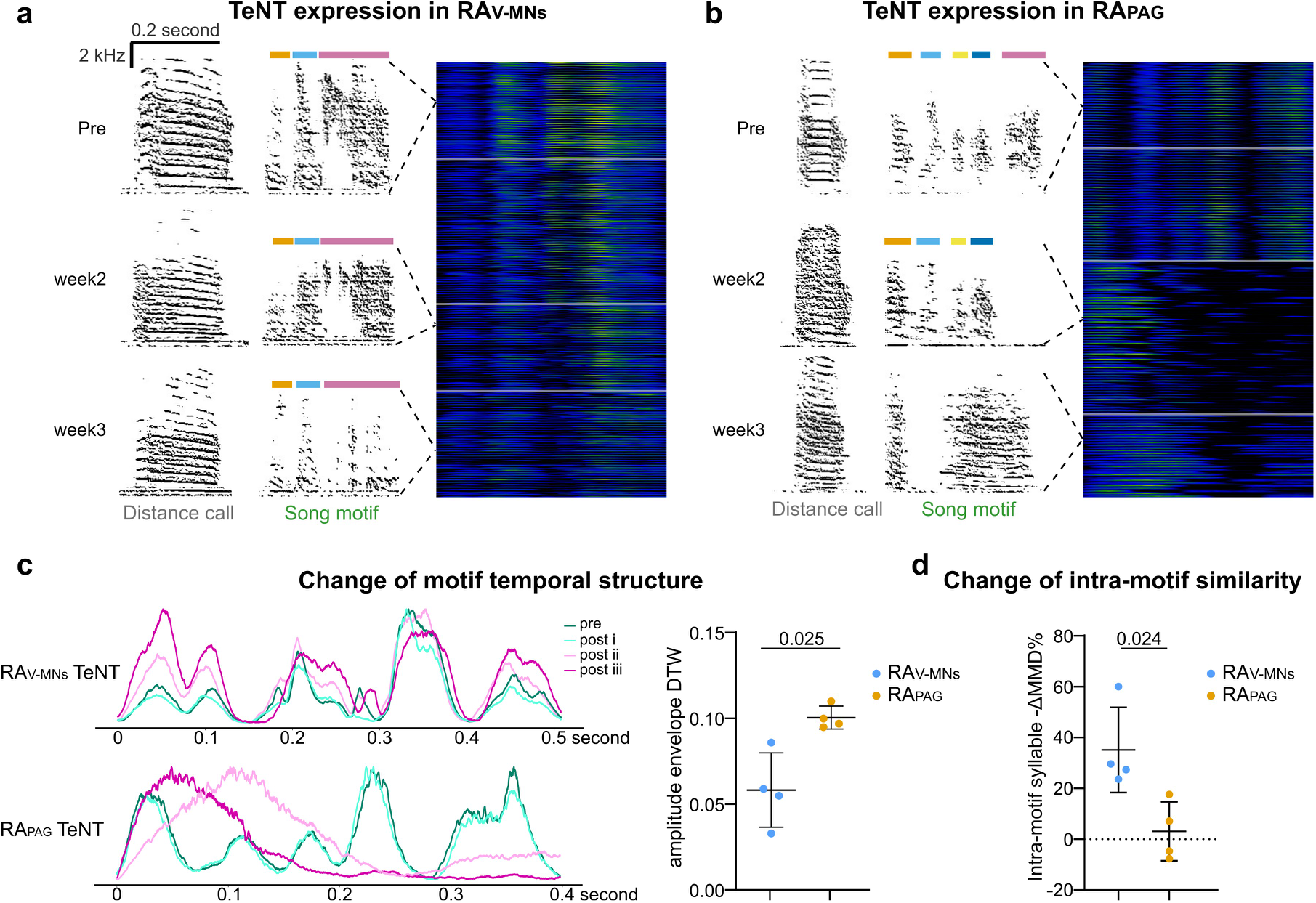
Pathways for controlling temporal and spectral features in learned vocalizations. **(a and b)** Expression TeNT in the two RA populations affected song production. Left, song spectrograms show change of song motifs and distance calls pre and post TeNT expression. Right, stacked motifs pre and week1,2,3 post TeNT viral injection. Colored bars mark each syllable of the motifs. n=4 animals for each group. **(c)** Change of motif temporal structure after TeNT expression. Left, motif amplitude envelopes pre and post viral injections. The normalized and averaged amplitude envelopes are calculated by averaging multiple (>10 motifs) amplitude envelopes from the same viral expression stage. Different colors indicate different viral expression stages. Right, Dynamic time warping (DTW) distance between averaged amplitude envelopes of the pre and post iii stage. Welch’s t test, comparing two TeNT groups. **(d)** Change of intra-motif similarity. Difference of syllables within the same motifs is calculated through Maximum Mean Discrepancy (MMD) and negative intra-motif MMD change before and after TeNT expression as intra-motif similarity change. Welch’s t test.

Silencing either pathway disrupted song but did so in observably different ways. To quantitatively examine these disruptions, we first plotted the normalized amplitude envelope of songs before and after TeNT expression. Dynamic time warping revealed that silencing the RA_V-MNs_ pathway had only minor impact on the timing of song syllables, while silencing the RA_PAG_ pathway severely degraded the temporal structure of learned songs via a progressive loss of song syllables and the inability to complete a song motif (**Fig. 2a-c, and Extended Data Fig. 2b,c**). We found that TeNT expression in either palliobulbar pathway caused disruptions to the spectral features that characterize individual syllables. We reasoned that if birds lost the ability to control the spectral content of their vocalizations that their song syllables would become more similar to one another. To examine this, we compared how individual syllables within a motif differ from one another by measuring intra-motif similarity using a deep convolutional neural network trained with over 16,000 manually annotated syllables from 21 adult zebra finches and calculating the Maximum Mean Discrepancy (MMD) between syllables^53^. This revealed that TeNT expression in RA_V-MNs_ results in syllables losing their individual qualities significantly more than when we express TeNT in the RA_PAG_ pathway (**Fig. 2d and Extended Data Fig. 2b,c and e**).

Together, results of these genetic silencing experiments indicate that the RA_PAG_ pathway is essential for producing the learned temporal patterns of song (i.e., the timing of syllable onsets and offsets). In contrast, RA_V-MNs_ control the learned spectral features within the syllables. Considering the potential bioacoustic mechanisms, our experiments support a model in which the direct RA-V-MNs pathway regulates dexterous syringeal and respiratory movements needed for producing the learned spectral structures in song syllables, whereas the RA-PAG pathway regulates a phonatory brainstem circuit that instructs vocal-respiratory timing with the millisecond precision needed for controlling the learned temporal structure of song.

### Functional and anatomical organization of the PAG

GABAergic projections from the limbic system have been shown to disinhibit the PAG and release production of innate vocal repertoires in mice^38^. Nevertheless, how excitatory palliobulbar projections interact with the PAG to control vocal-respiratory timing is not known. As a starting point to address this question we used anterograde viral jumping to label PAG neurons receiving synaptic input from RA. As this approach is not commonly used in the songbird^54^, we first verified this strategy by labeling V-MNs in the medulla through viral jumping from RA axons and Cre recombinase dependent expression of mScarlet in post-synaptic neurons (**Extended Data Fig. 3a**; n = 3 birds and 3 hemispheres). This revealed post-synaptic neuronal labelling patterns consistent with the presynaptic labeling of RA axons (**Fig. 1e**). We next labeled PAG neurons receiving synaptic input from RA and used RNA fluorescent in situ hybridization (FISH) to label glutamatergic and GABAergic neuronal populations (**Fig. 3a**). This revealed that RA predominantly innervates excitatory neurons in the PAG. Consistent with the stark effect of RA_PAG_ silencing, this finding helps rule out a simple scenario in which RA inhibits the PAG via a widespread projection onto local inhibitory neurons. Rather, it suggests that RA excites PAG and that the PAG may help actively control song.

**Fig. 3.**
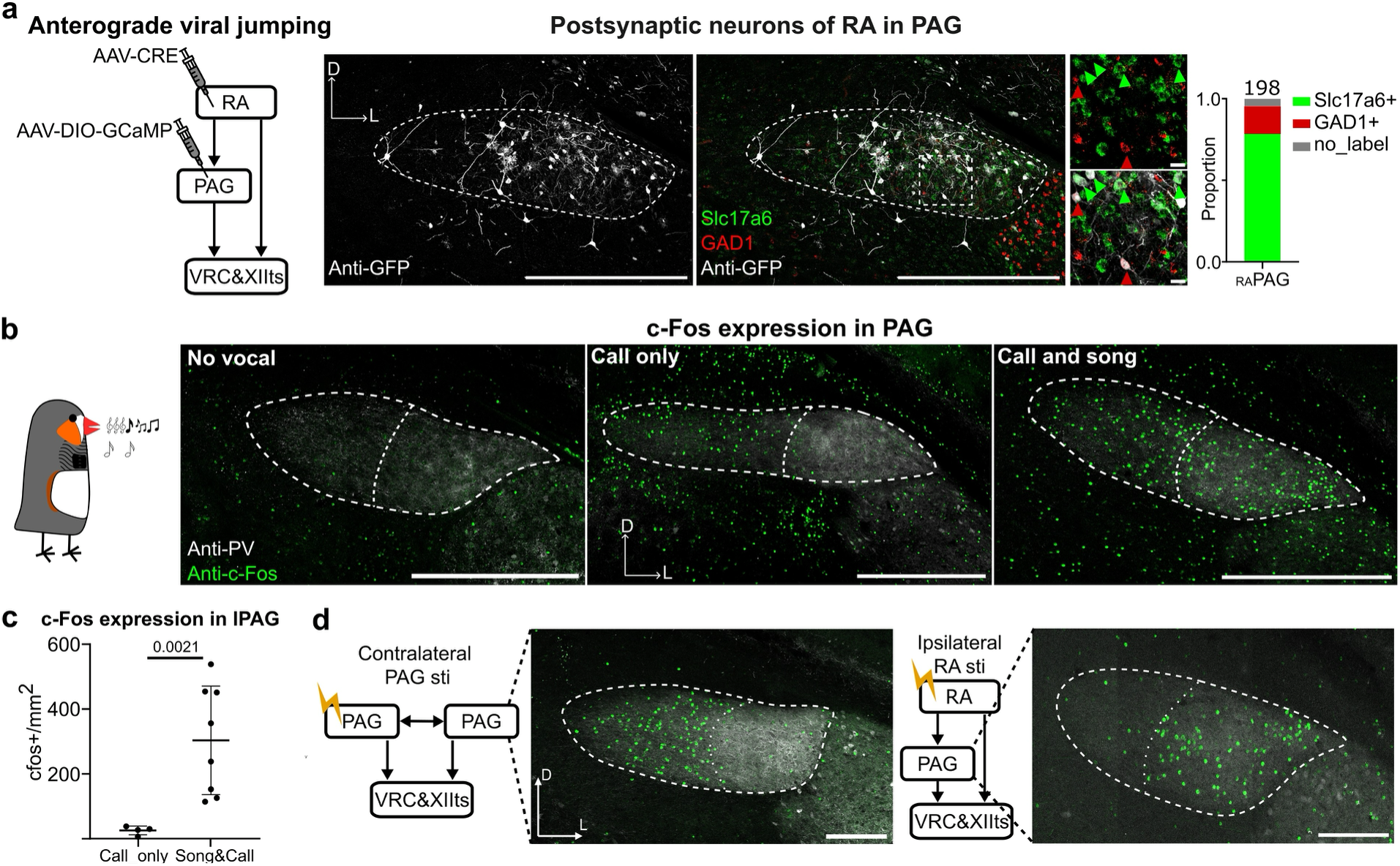
Functional organization of zebra finch PAG. **(a)** Both glutamatergic and GABAergic PAG neurons receive RA input. Left, the viral-injection schematic of anterograde viral jumping. Middle, PAG neurons are labelled by anti-GFP (grey), FISH-Slc17a6 (green) and FISH-GAD1 (red). Dash contours outline PAG, and the dashed box marks zoom-in regions. Arrowheads (red or green) point out glutamatergic or GABAergic neurons co-labeled with anti-GFP. Right, Slc17a6/GAD1 ratio is quantified from 198 neurons (from 3 animals). Scale bars, 500 μm, except the zoom-in images (20 μm). **(b)** c-Fos expression in PAG after different vocal activities. Three groups of animals are included, non-vocalizing birds, birds that only produced calls, and birds that produced songs and calls. PAG is highlighted by anti-PV signal in grey. Anti-c-Fos signal is in green. Dashed contours outline the medial and lateral port of the vocal PAG. Scale bar, 500 μm. **(c)** c-Fos expression in the lateral PAG after only calling and both singing and calling. n=4 for call only group, n=8 for song and call group. Welch’s t test. **(d)** Electrical stimulation induced c-Fos expression in the vocal PAG. Circuit schematic showing electrical stimulation of contralateral PAG (left) and ipsilateral RA (right). Dashed contours and lines outline the medial and lateral port of PAG. Scale bar, 200 μm.

The mammalian PAG is organized into molecular and functional subregions and electrical stimulation of different locations gate production of distinct innate vocalizations^37,55–57^. How the songbird vocal PAG might be anatomically and functionally organized to control calls and song is not known. Through a series of anatomical tracing, functional, and molecular mapping experiments, we found that the songbird vocal PAG is organized into lateral and medial subregions which are respectively associated with learned and innate vocalizations.

First, we found that V-MNs projecting neurons in PAG (PAG_V-MNs_) are scattered throughout the vocal PAG but are denser in its more lateral aspect (**Extended Data Fig. 3b,c**). We find these neurons are excitatory and characterized by their expression of the glutamate receptor subunit GluR-5 (GRIK1^+^) (**Extended Data Fig. 3c**)^58^. A non-overlapping class of mostly GRIK1 negative projection neurons, which innervate the contralateral PAG (PAG_PAG_), are found in the medial vocal PAG. Inputs from RA are also well organized (**Extended Data Fig. 3d,e**). Although RA innervates throughout the vocal PAG, its axon terminals are denser in the lateral PAG and only in the lateral PAG do they form basket like terminals suggesting strong synaptic input onto PAG neurons (**Extended Data Fig. 3d**). RA axons are parvalbumin (PV) positive^59^ and PV staining supports a significant medial lateral gradient within the PAG (**Extended Data Fig. 3e**), which is eliminated following RA lesions (**Extended Data Fig. 3f)**. Thus, we find that the PAG projection to the V-MNs and contralateral PAG neurons are topographically organized, and RA’s inputs are anatomically well positioned to influence the lateral PAG_V-MNs_ pathway.

To examine whether and how the medial-lateral subregions of PAG are involved in vocal motor control, we mapped the activity dependent expression of the immediate early gene c-Fos after production of calls or production of calls and song (**Fig. 3b**)^60,61^. We anatomically defined the medial and lateral aspects of the PAG based on RA innervation patterns, as shown through anti-PV antibody labeling (**Extended Data Fig. 3d-g**). Negligible c-Fos was detected in PAG if birds did not produce any vocalizations (**Fig. 3b and Extended Data Fig. 3h**). Interestingly, only the medial PAG was c-Fos positive following production of calls (**Fig. 3b,c**). Although it is not possible to have birds produce songs without interleaved calls, production of songs and calls resulted in c-Fos expression in the medial and lateral portions of the vocal PAG. Further, the amount of c-Fos expression in the lateral PAG was strongly correlated with the number of song motifs, but not with the number of calls generated (**Extended Data Fig. 3i**).

Given that input pathways from the contralateral PAG and ipsilateral RA are topographically organized along the medial lateral axis of the PAG, we wondered if our c-Fos labeling directly related to these separate input pathways. To examine this, we electrically stimulated either the contralateral PAG or the ipsilateral RA (most RA axons do not decussate) in anesthetized birds to examine if we could drive spatially restricted c-Fos expression in the PAG (**Fig. 3d**). We found that electrical stimulation of the contralateral PAG resulted in strong c-Fos expression only in the medial PAG. In contrast, electrical stimulation of the ipsilateral RA resulted in strong c-Fos expression in both the lateral and medial PAG (**Extended Data Fig. 3j**). Together, these findings indicate that the PAG and RA inputs are functionally and structurally organized into subregions supporting innate and learned vocalizations.

### PAG uniquely encodes learned and innate vocalizations

It is not known how activity in the mammalian or avian PAG may encode learned vocalizations. To further investigate the mechanisms of descending vocal motor control, we made high-density Neuropixels 2.0 neural recordings from the PAG in four freely behaving male zebra finches (**Fig. 4a and Extended Data Fig. 4**). We found that units within the PAG reliably modulate their firing rates during production of songs and calls (**Fig. 4b and Extended Data Fig. 5a-c**). Fitting with our anatomical descriptions of PAG organization, we also found that the lateral portion of the vocal PAG showed a higher firing rate during songs than the medial PAG (**Extended Data Fig. 4b**).

**Fig. 4.**
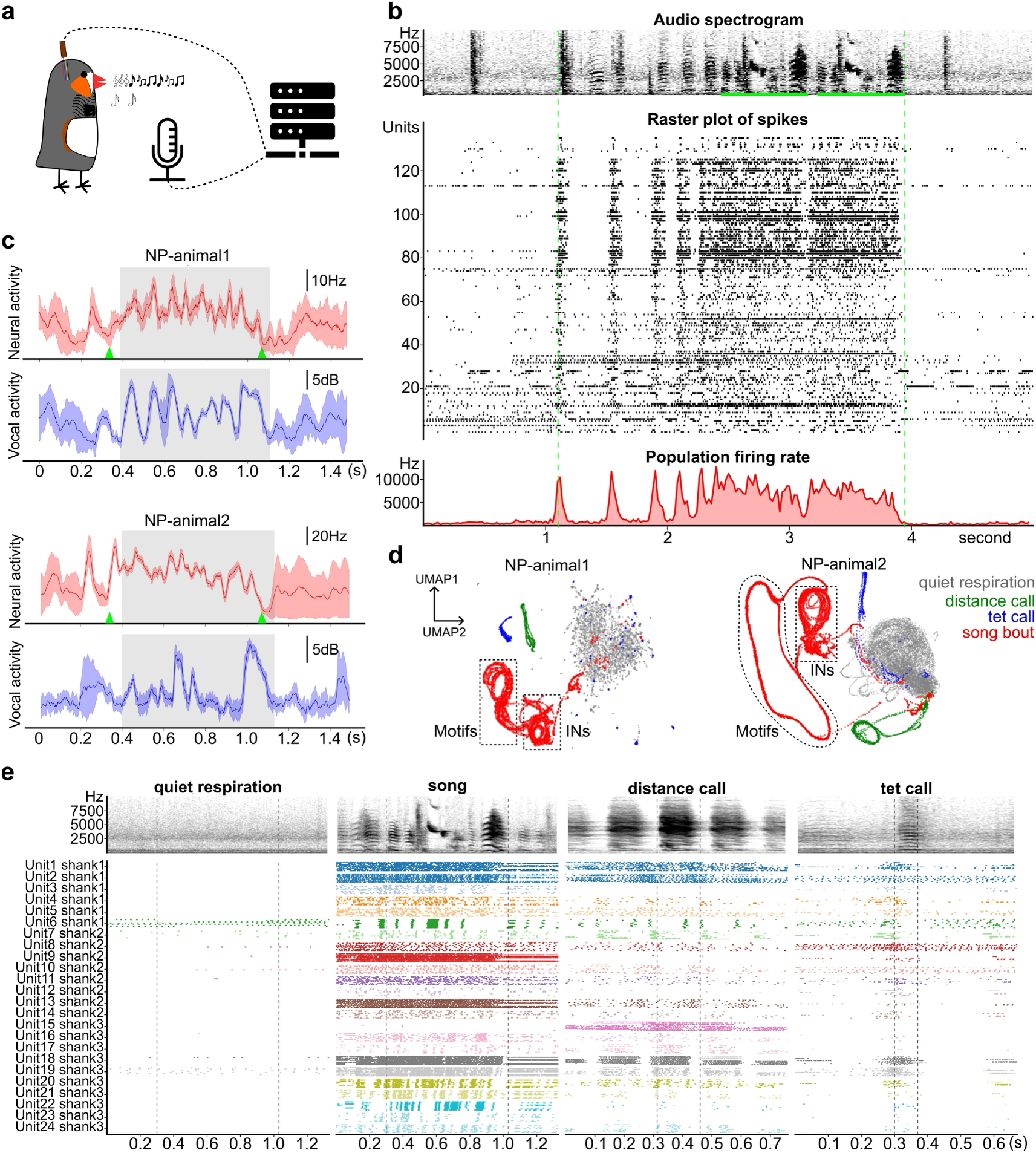
The PAG uniquely encodes songs and calls. **(a)** Schematic of experimental setup to collect Neuropixels data recordings in freely singing animals. **(b)** Synchronized song spectrogram and raster plot of vocal-related unit spikes. Top, song spectrogram, in which song motif renditions are underlined by green. Medial, raster plot of unit spikes. Each row is an isolated unit from Neuropixels recording. Each black dot is a spike event of the units. Bottom, Summed firing rate across the population. The vertical dashed green lines indicate the beginning of the first introductory note and end of the song bout. **(c)** Stereotyped song motif and unit activity of two recorded animals. The red and blue lines show curves of population firing pattern and vocal amplitude, averaged from n=8 and n=9 motif renditions, respectively. The shadow outlines their standard deviation. Green arrowheads identify onsets and offsets of neural modulation. **(d)** Trajectory UMAP of unit activity during different vocal states. Population spike patterns in different vocal events are segmented into overlapping windows which are embedded into the UMAP space. Dashed contours highlight time windows of introductory notes and motifs within song bouts. **(e)** Firing patterns of well isolated somatic units during different behaviors. Audio spectrograms are generated by averaging vocal renditions. Raster plot illustrates the stacked firing activity of each unit while the animal was producing the same vocalizations. Firing dots of each unit are colored uniquely and synchronized to the onset of the vocalization, signified by the first grey dashed line in each panel.

We identified 311 vocal related units from our PAG recordings (**Fig. 4b and Extended Data Fig. 4 and 5a**). We found that the activity of these units was highly stereotyped during song and preceded vocal onsets by ∼30ms (**Fig. 4c and Extended Data Fig. 7a)**, indicating that the PAG plays a premotor role in the control of song syllables and calls. We next examined if population activity in PAG uniquely encodes these behaviors. Using dimensionality reduction approaches we plotted the trajectory of neural activity from each bird in latent space during quiet respiration, production of song motifs, distance calls, and during tet calls (**Fig. 4d**). This revealed that each of these behavioral states is encoded by unique, non-overlapping trajectories. Thus, premotor population activity across the PAG differentially encodes the learned song and different call types in the zebra finch repertoire.

To explore the premotor activity patterns of well-isolated, simultaneously recorded somatic units, we aligned neuronal activity to vocal onsets (**Fig. 4e and Extended Data Fig. 4c-f, 5d**). Most somatic units increased firing rates during vocalizations (**Extended Data Fig. 5b**), and individual units exhibited different levels of activity during learned song versus innate calls (**Extended Data Fig. 5c**). PAG single units showed remarkably precise and stereotyped bursting activity during songs and different call types, and across the population, individual units had distinctive firing patterns. Most neurons were active during calls and song (40/60 units), some were active selectively during songs (12/60 units) while others were only active during one call type or another (2/60 units for distance calls; 6/60 units for tet calls). Together, these data indicate that PAG neurons directly encode both songs and calls using precise and reliable firing patterns.

### PAG is required for learned and innate vocalizations

To directly test the role of the PAG in song and calls, we selectively silenced the PAG_V-MNs_ pathway using Cre-dependent TeNT (**Fig. 5a**; n = 9 birds). We found that birds’ songs and calls began to rapidly degrade 7-10 days post intracranial viral injections, a timeframe consistent with AAV-TeNT expression in the rodent PAG^39,62^ (**Fig. 5b**). After two weeks of viral expression, the number of song bouts and total vocal activity, including all innate calls, dropped significantly in TeNT birds but not in controls, which expressed only Cre-dependent mScarlet fluorescent protein. We found that several TeNT birds were completely muted of their songs and calls. This strong phenotype correlated with higher levels and duration of viral expression in the PAG_V-MNs_ pathway, providing further confidence that effects on vocal production were directly related to silencing of specific projection neurons in the PAG (**Fig. 5c and Extended Data Fig. 6a,b**). Together, these findings reveal that the PAG_V-MNs_ pathway is required for production of learned and innate vocalizations.

**Fig. 5.**
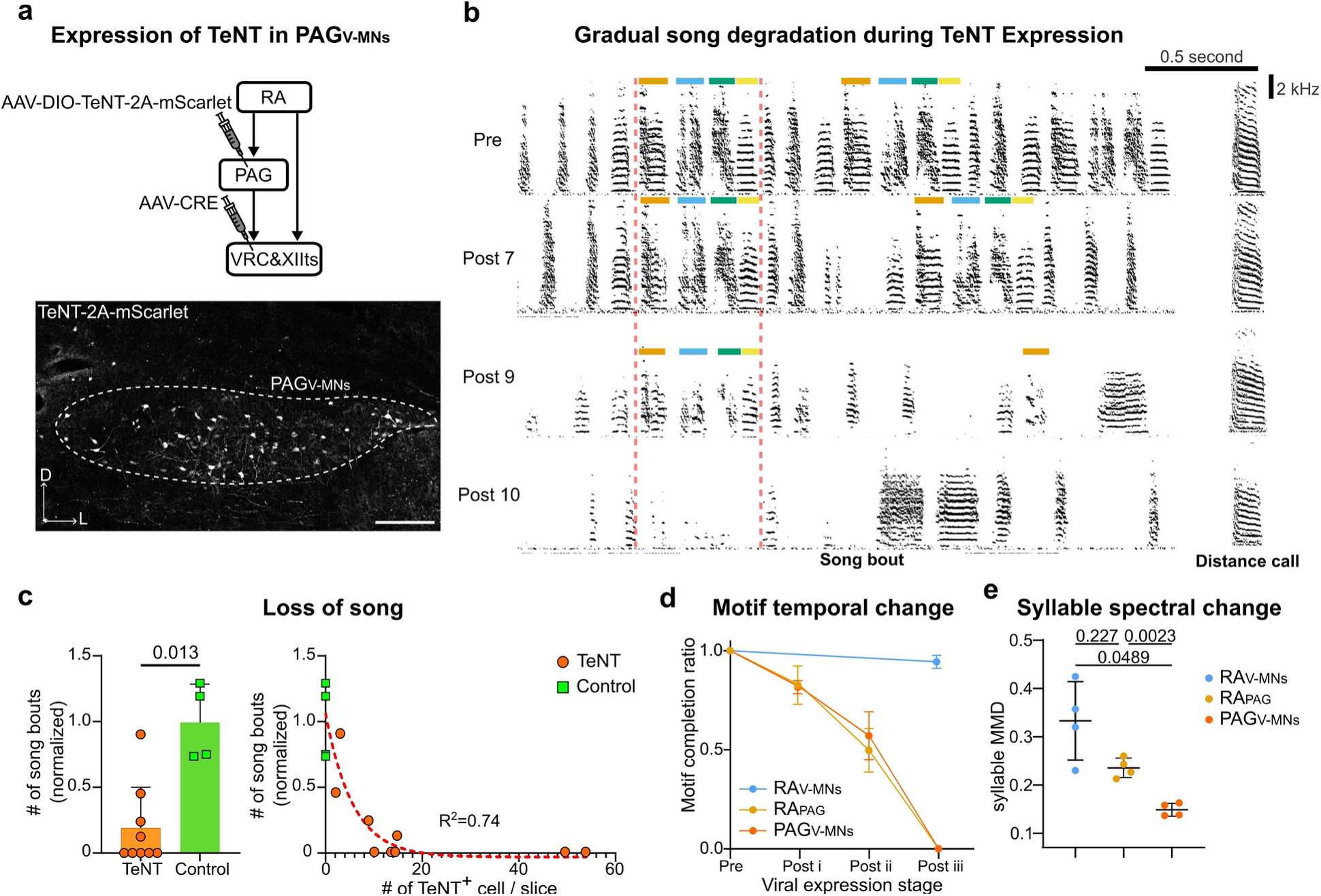
PAG is necessary for learned vocalization. **(a)** Expressing TeNT in PAG_V-MNs_. Top, the schematic of viral injection to specifically express TeNT in PAG_V-MNs_. Bottom, image showing TeNT expression in PAG_V-MNs_. The dashed contour outlines the vocal PAG. Scale bar, 200 μm. **(b)** Song spectrogram pre and post the viral injection. The same song syllable is highlighted by the same color across the viral expression timeline. The vertical red dashed lines outline motifs. **(c)** Relative number of song bouts produced after TeNT expression. Normalized song number is calculated by number of songs produced in post day14 divided by the number a day before viral injection. Left, a group plot of TeNT and control groups. Mann Whitney test. Right, correlation of normalized songs and viral expression levels in the TeNT groups. A one-phase decay curve was fitted to the TeNT group data. **(d)** Motif completion ratio pre and post TeNT expression. The ratio is quantified by the number of syllables finished (once initiating the first syllable) divided by total syllable number. The timeline of PAG and RA TeNT group is aligned by designating the last singing day of PAG lesion cohort and the second week of RA lesion cohort as Post ii. n=4 for each group. **(e)** Change of syllables similarity. Brown-Forsythe and Welch ANOVA tests and Dunnett’s T3 multiple comparisons test.

Before birds became mute, we observed a progressive loss of song syllables and the inability to complete a song motif. This behavioral effect closely matched the effects of silencing the RA_PAG_ pathway (**Fig. 5d**). However, unlike the PAG_V-MNs_ manipulations, silencing RA_PAG_ pathway did not mute birds as they were still able to produce calls and introductory notes. Also, TeNT expression in PAG_V-MNs_ neurons caused a minor spectral change in song syllables and resulted in weaker disruption of spectral structure than seen when silencing either of the palliobulbar pathways (**Fig. 5e**). Because PAG is thought to gate production of innate vocalizations, we were surprised to find that TeNT expression in either RA pathway also disrupted spectral properties of calls more than TeNT expression in PAG_V-MNs_ (**Extended Data Fig. 6c**). These findings suggest that the fine spectral features of male specific calls may be regulated through a similar circuit mechanism that controls song syllables.

Lastly, we noticed that the amplitude of song syllables and calls gradually declined before birds completely lost the ability to produce vocalizations (**Extended Data Fig. 6d,e**). Silencing either RA_PAG_ or RA_V-MNs_ caused a similar decrease in the amplitude of song syllables (**Extended Data Fig. 6f**). Yet, only silencing the RA_PAG_ neurons, reduced the amplitude of calls. This suggests that this palliobulbar pathway helps control vocal respiratory pressure needed for phonation. This would align well with research in mammals showing that increasing electrical stimulation intensity in the PAG increases the amplitude of the evoked vocalizations^63^. To test this in zebra finches, we elicited vocalizations in anesthetized birds using electrical stimulation in the PAG (**Extended Data Fig. 6g**). We found electrical stimulation evoked call-like harmonic vocalizations with short latencies, consistent with previous studies^64,65^, and that increasing current intensity evoked higher amplitude vocalizations. These results further support a model in which the RA-PAG-V-MNs pathway regulates vocal-respiratory timing and the vigor underlying production of learned vocalizations.

### Hierarchical vocal timing codes in RA and PAG

To examine the RA-PAG-V-MNs pathway in more detail, we asked how RA and PAG encode for the different phases of vocal-respiratory timing. In two of the birds in which we obtained Neuropixels recordings from the PAG, we were able to subsequently also make high-density recordings from RA during singing (**Fig. 6a,b**). In these birds we found that RA singing related activity led PAG by 2–3 milliseconds, supporting the view of top-down motor control of respiratory timing.

**Fig. 6.**
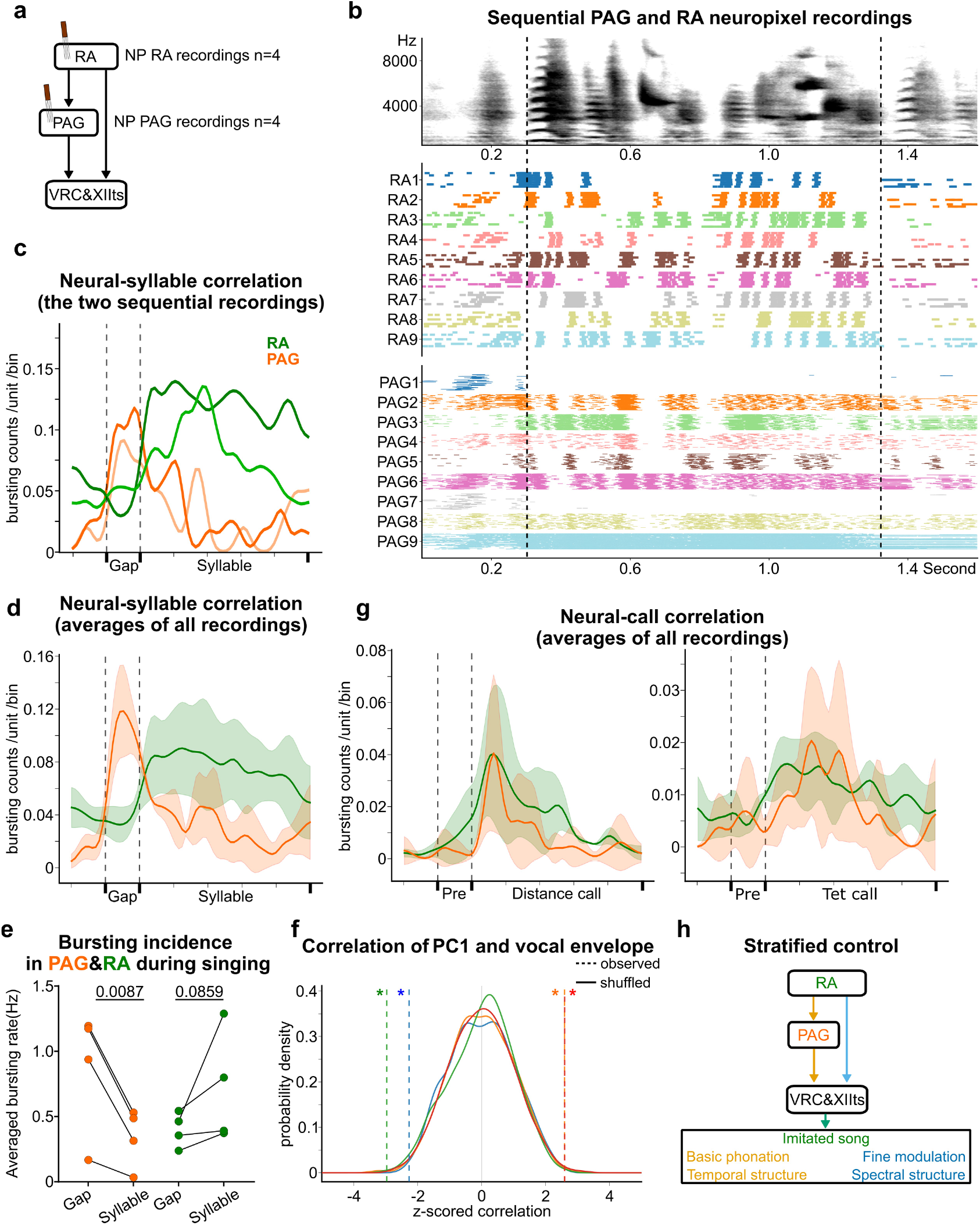
Vocal timing codes in RA and PAG. **(a)** Schematic of Neuropixel recordings in the descending vocal circuits. **(b)** Example of Neuropixel recordings in RA and PAG during singing in the same zebra finch. Raster plots of RA and PAG units aligned with the birds stereotyped song. **(c)** Neural activity of RA and PAG aligned with vocal onsets and offsets. Bursting activity of well isolated somatic units of RA and PAG was time-shifted and segmented into pre-gap, gap and syllable periods, then time-warped linearly(pre:gap:syllable=1:1:5). Lighter and darker colors indicate the two different birds in which we recorded from both PAG and RA. **(d)** Averaged bursting curves aligned with syllable timing, including two animals of panel C and four additional animals with recordings from either RA or PAG. **(e)** Quantification of bursting incidence during syllables and gaps. Ratio paired t test. **(f)** Z-scored pearson correlation bewteen principal component 1 of RA population activity and vocal amplitude envelopes. Z-scores and significance were calculated based on 1000 burst-shifting shuffles of each animal. Solid curves are distribution probability curves of shuffled correlation values, and vertical dash lines are the observed correlation value of each animal. **(g)** Neural activity of RA and PAG during distance calls and tet calls. **(h)** Stratified control model. RA regulates song production (green). RA-PAG-V-MNs pathway regulates basic phonation for temporal structure of learned vocalization (orange), while direct RA-V-MNs pathway dexterously modulates the fine spectral structure (blue).

At the single neuron level, PAG neurons are found to be positively or negatively correlated with the song vocal amplitude envelope (**Extended Data Fig. 7b**). Overall correlation coefficients trend to be more negative when comparing all PAG neurons with RA neurons (**Extended Data Fig. 7c**). This suggests that PAG units are tuned by vocal-respiration timing and may better encode the silent gaps between song syllables. To explore temporal coding at the population level, we used linear time warping to align bursting activity in RA and PAG associated with individual song syllables and their preceding silent gaps - gestures typically associated with expiration and inspiration (**Fig. 6b,c**). This revealed that RA bursting activity aligns with expiratory phases, while bursting in PAG peaks with the inspiratory phases preceding syllable onsets. Recordings from RA in two additional birds combined with analysis of our other PAG recordings, confirmed our initial impressions of burst timing (**Fig. 6d**). Indeed, we found that the rate of bursting in PAG was significantly higher during silent gaps than syllables, while gap-syllable bursting in RA trended in the opposite direction (**Fig. 6e**). Simultaneously recording hundreds of RA neurons also allowed us to extract the main population vector through principal component analysis. This revealed that PC1 of RA significantly correlated with the song vocal amplitude envelope (**Fig. 6f**).

Given the established and evolutionarily conserved role of the PAG in motor control and gating of innate vocalizations^39,49^, we wondered whether this held for zebra finch distance and tet calls. Interestingly, bursting activity in PAG and RA both peaked during production of calls (**Fig. 6g)**.

This is consistent with a premotor role of the PAG and RA in the control of innate vocalizations and further suggests that inspiratory-related timing of PAG activity during song is an adaptive specialization associated with the precise temporal structure of learned song syllables. Altogether our findings suggest stratified control of vocalizations via palliobulbar pathways (**Fig. 6h**)^3,46,66^. The largely non-overlapping palliobulbar projections from RA to the PAG and V-MNs respectively control the timing and spectral features of learned song. The PAG is a final common pathway for producing any vocalization, it uniquely encodes learned and innate vocalizations, and functions critically in the control of vocal timing.

## DISCUSSION

The underlying circuit mechanisms for the control of learned vocalizations have been poorly understood. We show that two major palliobulbar pathways from RA control different aspects of learned birdsong. The pallio-PAG pathway regulates the temporal structure of song syllables, while the pallio-medullary pathway controls their fine acoustic properties (**Fig. 2**). Moreover, we find an anatomical and functional organization across medial lateral aspects of the PAG (**Fig. 3**). Through electrophysiological recordings, we show that activity in the PAG uniquely encodes the song motif and different call types (**Fig. 4**). We also find that functionally silencing PAG inputs to the medulla can completely mute birds (**Fig. 5**). Although difficult to interpret, strokes impacting the PAG have been reported to cause mutism in people^42,67,68^, suggesting convergent circuit mechanisms in the evolution of birdsong and speech motor control. Additionally, we find that silencing palliobulbar projections to either PAG or medulla has a stronger effect on the acoustics of calls than silencing PAG_V-MNs_. This suggests overlapping circuit mechanisms in control of calls and syllables^32^, a finding that could have important implications for species with various levels of vocal flexibility. Lastly, we explore vocal temporal codes in RA and PAG and find the PAG is preferentially active during silent gaps between song syllables (**Fig. 6**), which suggests that the precise temporal control of the learned songs is achieved by regulating inspiratory timing via the RA-PAG pathway.

Studies in mammals demonstrate that different regions of PAG may regulate different types of innate vocalizations^55,69^. In zebra finches, we show that PAG neurons are molecularly and anatomically organized. The medial and lateral subregions are differentially involved in innate and learned vocalizations. These findings suggest that partially overlapping PAG neuronal populations may regulate different aspects of vocalizations. Neurons in the medial PAG tended to be active during all vocalizations and contained interhemispheric PAG_PAG_ neurons. Thus, the medial PAG may mediate bilateral coordination and expiratory drive, while PAG neurons in more lateral aspects are biased to song and may encode identity of each type of vocalization. In the case of learned songs, we show that PAG_V-MNs_ are necessary for phonation. However, we cannot rule out the possibility that the PAG pathway also regulates some other aspects of learned vocalizations. For example, PAG could encode a genetically inherited prototype of syllables or gate motor prototypes in hindbrain circuits which are further refined by the direct RA-medullary pathways during learning^70,71^.

Our findings also provide potential insights into the circuit mechanisms for vocal production in other vocal learning species. We show that the PAG to V-MNs pathway is obligatory for production of innate and learned vocalizations alike. In humans, the motor cortex is thought to innervate laryngeal motoneurons and other V-MNs to facilitate speech production, analogous to the RA_V-MNs_ pathway^1,3^. Some models emphasize the seemingly exclusive role of this pathway in production of speech^4,47,72,73^, while others have emphasized the integrated role of this pathway and the PAG related circuits in speech^3,66^. Studies in non-human primates show that ACC is involved in and may be able to initiate vocalizations through its projections to the PAG^55,74–76^. Given the currently known anatomy, our results support the view that human speech may engage the PAG and that descending projections from the ACC help control respiratory patterns while descending circuits from the motor cortex control the larynx and articulation. A similar collaboration of volitional motor circuits and emotional circuits is also reported in orofacial behaviors of nonhuman primates^77,78^, suggesting a common theme in how the brain balances the organization of circuits underlying dexterous motor behaviors. Thus, the distinct functions of the direct and indirect, via PAG, projections to the medulla suggest an evolutionary principle for motor control.

In summary, this research reveals how palliobulbar projections can have separable functions in the control of a learned vocal behavior. We further reveal the essential function of the PAG in producing learned song and integrating both innate and learned behaviors. Our results therefore provide unique insights into how palliobulbar pathways control learned communicative behaviors, with potential implications for production of learned behaviors in other taxa.

## Acknowledgments

We thank members of the Roberts lab for comments on an initial version of the manuscript, Dr. Jie Cao for cloning of all viral constructs used in this research. We thank Dr. Wei Xu for providing TeNT and mScarlet-2A-syb2-eGFP plasmids, Dr. Brad Pfeiffer for sharing 3D printers, Jennifer Holdway, Luis Garcia, Jeremy Brusch and Rico Cabuco for laboratory support. We thank Jesse Hilton, Richonda Hunte, and Pamela Jennings for administrative support.

## Funding

We thank the National Institutes of Health for supporting this research through grants.

National Institutes of Health grant UF1NS115821 (TFR)

National Institutes of Health grant R01DC020333 (TFR)

National Institutes of Health grant R01NS134574 (TFR)

National Institutes of Health grant R01NS108424 (TFR)

## Author contributions

Conceptualization: ZZ, TFR

Methodology: ZZ

Investigation: ZZ

Visualization: ZZ, TFR

Funding acquisition: TFR

Project administration: TFR

Supervision: TFR

Writing – original draft: ZZ

Writing – review & editing: ZZ, TFR

## Competing interests

Authors declare no competing interests.

## Data, code, and materials availability

The datasets generated during and/or analyzed during the current study will be made publicly available in the Texas Data repository upon publication. Analysis codes used in this research will be made available on GitHub upon publication.

## Materials and Methods

### Animals

Adult (100-200 days post hatch) male zebra finches were used in this study. Birds were kept under a 14:10-h light:dark cycle with free access to water and food in a group-housed colony cage or a sound-isolated chamber for audio recording. All experiments were performed under protocols approved by Animal Care and Use Committee at UT Southwestern Medical Center.

### Viral vectors

Custom-made adeno-associated viruses were used in the experiments. Vectors include: rAAV2/9-CAG-cDIO-TeNT-2A-mScarlet, rAAV2/9-CMV-Cre-eGFP, rAAV2/9-hSyn1-mScarlet-2A-syb2-eGFP, rAAV2/9-CBH-fDIO-mScarlet-2A-syb2-V5, rAAV2/9-CBH-Flp, rAAV2/9-CAG-cDIO-GCaMP8m, rAAV2/9-CAG-cDIO-mScarlet.

### Brain surgery

Birds were anaesthetized by 1% isoflurane and head-fixed on custom stereotaxic rig. Body temperature and breathing were monitored during surgeries. RA and hindbrain vocal circuits (VRC and XIIts) were located by electrophysiologic mapping assisted by previous coordinates relative to the bifurcation of the sagittal sinus. The PAG was identified by electrical stimulation (2s 10-30 µA 100Hz square wave). Actual coordinates varied across animals (head angle, lateral, anterior-posterior, depth (in mm)): for RA (20°, 2.0-2.7, 1.0-2.0, 1.7-2.5); for PAG, (45°-55°, 2.4, 1.0-2.0, 4.7-6.2); for XIIts and VRC (45°-55°, 0.3 and 1.0, 2.0-3.0, 5.0-5.7).

Stereotaxic injections were performed using previously described procedures^21,79,80^. 1.5-2 µl viral solution was injected at 1 nl/s into each brain regions, 150 µl chemical tracers (Dextran-Alexa488, Invitrogen; FastBlue, Polisciences; CTB-Alexa647, Invitrogen) or pharmacological drug (a 1:1 mix of 1% ibotenic acid and 100mM Quisqualic acid) 150nl was injected at 33nl/s into each brain region.

### Audio recording and analysis

Vocalizations were recorded using Sound Analysis Pro (SAP2011), except vocalizations during electrical stimulation and Neuropixels recording, which were recorded by Audacity and Open Ephys, respectively. Recorded song files were first cleaned by manually deleting pure noise. Song bouts were segmented based on amplitude threshold, and motifs and specific male calls are segmented based on template matching of amplitude envelope or manually curated. All the segmented audio files were further validated and cleaned manually.

Amplitude of syllables and calls was calculated using AVN^81^ acoustics module. Normalized amplitude envelopes were generated by averaging multiple motifs aligned at the onset of the first syllable or the last introductory note and normalized by the max value. Dynamic time warping was used to quantify similarity between amplitude envelopes. Motif completion ratio was quantified by manually counting how many syllables were produced in the motif once the first syllables presented. Acoustic similarity between syllables was calculated by AVN’s triplet-loss model combined with Maximum Mean Discrepancy (MMD)^53^. Syllables were embedded in the latent spaces of a pre-trained triplet-loss model. Syllable embedding was visualized through UMAP.

MMD was calculated to quantify the distance of syllables in the latent spaces. Intra-motif similarity change was calculated by negative change of average MMD between different syllables of the core motif. The call correlation heatmap was generated through calculating Pearson correlation of the segments of their spectrogram before and after TeNT expression^17,19^. Call similarity was calculated by averaging the diagonal values in the correlation heatmap. All analyses were performed using Python code and GraphPad, except where explicitly stated otherwise.

Animals were single housed in a recording chamber for at least a week. PAG was located with electrical stimulation, then a multi-shank Neuropixels 2.0 probe was implanted into the brain region where electrical stimulation induced calls (NP-animal1,2,3,4 with 2,3,1,1 shanks in PAG, respectively). RA was located by standard coordinates and electrophysiologic mapping. We used the ‘indie’ (independent dovetail implants for electrophysiology) fixture^82^ to hold the probe and it was secured with super glue gel and dental cement. The indie probe fixture was connected to a custom counterweight system. The day after implantation, a female was presented to elicit vocalizations. Data collection was confined to the 2 days following probe implantation.

We used the OneBox system with Open Ephys for data acquisition. An external variable TTL signal was generated by a computer, amplified and bifurcated into a sync input and an ADC recording channel of OneBox. 384 channels were simultaneously recorded with the external TTL signal at 30,000 Hz. Audio recordings were obtained through another ADC channel in OneBox at 30,300.5 Hz.

Raw data were spike-sorted with Kilosort4.0 using default parameters. Sorted units were validated through Phy2. Custom python codes were used to synchronize spike data and audio data via alignment with TTL signals. The plots of Neuropixels recordings were analyzed and generated using custom python codes. Vocal related units were manually selected based on their firing pattern during vocalization. Trajectory UMAPs were generated with all the vocal-related units. Recording periods were separated into the four groups, tet calls, distance calls, song bouts, and quiet respiration (using the same duration as the song bouts). These periods were further divided into overlapping time windows, optimized for each animal (NP-animal1: 200 ms window, 3 ms step; NP-animal2: 100 ms window, 3 ms step). Each time window was further divided into five segments (40ms for NP-animal1, 20ms for NP-animal2) and the number of spikes during the 5 segments was counted for each unit. The overlapping windows (five segments X N number of units) were embedded into the UMAP space using custom Python code.

Population analyses included all the sorted vocal units. Further analyses on single units only included the well-isolated somatic units, defined by somatic spike waveform and low ISI violation. Time lapse between vocal activity and unit activity were calculated by sliding of neural “self”-correlation matrix to match with vocal “self”-correlation matrix^17^. Neural “self”-correlation matrix was generated by calculating Pearson correlations between unit activity during one motif and another motif (for example, there are 8 motifs from NP-animal1, so 36 neural correlation matrices were generated). Corresponding vocal motif-to-motif correlation matrices were generated. Then we shifted neural matrices forward or backward in the -50ms to +50ms window, and calculated Pearson correlation between the shifted neural matrix and the corresponding motif matrix. The best shifting timepoint was found by identifying the highest correlation score between the two matrices.

Correlation analysis between unit activity and vocal timing patterns was performed after shifting forward the neural activity based on the time lapse. Bursting peak events were detected through firing frequency curves where peak activities above 250 Hz firing rate were detected and extracted. To average neural activity (bursting and firing) across different silent gaps and syllables, a linear time-warping was applied to syllables, preceding silent gaps and 20ms-windows before each silent gap with a final ratio of 5:1:1.

To capture the dominant modes of neural population activity of RA, we applied Principal Component Analysis (PCA). For each animal, we constructed a feature matrix of dimensions *T × N*, where *T* is the number of time points (1 ms resolution, 10 ms sliding window) and *N* is the number of recorded units. Each element is firing rate of one unit during that 10ms-window. The first principal component (PC1) was extracted for subsequent correlation analysis. We generated a null distribution by shuffling spike times while preserving burst statistics, where we detected burst periods using criteria of ≥5-spikes firing peak within a 10 ms window, segmented non-burst periods into variable bins (1-5 ms), randomly permuted the temporal order of all periods (bursts and non-burst bins). This procedure maintained the overall firing rate and burst characteristics while destroying the temporal sequence of neural events. For each one of 1000 shuffle iterations, we recomputed PC1 and recalculated the correlation with the amplitude envelope.

### Tissue collection

Brain tissue was collected 3 days after tracer injection and at least 14 days after viral injection. Animals were anesthetized with Euthasol (Virbac, TX, USA) and transcardially perfused with 4% paraformaldehyde (PFA). Brains were extracted, post-fixed in 4% PFA on a 4 °C shaker overnight and switched to 30% sucrose 4% PFA overnight. 40 µm coronal sections were cut using a cryostat and stored in PBS or in 4% PFA for fluorescence in situ hybridization.

### Immunohistochemistry

Sections were washed in PBS on a shaker for 10 minutes 3 times, then incubated with blocking buffer (3% bovine serum albumin, 0.3% Trition X-100 in PBS) for 15 minutes and transferred to primary antibody solutions (1:1000 diluted primary antibodies in blocking buffer) overnight at 4 °C (anti-GPF (Invitrogen), anti-Parvalbumin (Swant), anti-c-Fos (Santa Cruz Biotechnology), anti-V5 (Invitrogen). Sections were washed 3 times in PBS, then incubated with secondary antibodies at room temperature for 2 hours and washed 3 times in PBS. Sections were mounted and coverslipped with Fluoromount-G and imaged with a confocal microscope, LSM 880. Image analyses were conducted via ImageJ Fiji.

### Fluorescence in situ hybridization

FISH was conducted using v3 in situ hybridization chain reaction (Molecular Instruments). Probes were designed and produced by Molecular Instruments using the NCBI database (Slc17a6 NM_001309508, GAD1 XM_002198498, GRIK1 XM_030264380). Sections were post-fixed in 4% PFA for at least 15 minutes, washed in PBS 3 times, incubated in PBS with 5% SDS for 15 minutes and washed in 2×sodium chloride sodium citrate 0.1% Tween20 (2×SSCT) for 10 minutes 3 times. Sections were then incubated in the probe hybridization buffer (Molecular Instruments) for 10 minutes and then transferred to the probe solution (1% HCR probes in the hybridization buffer) at 37 °C for 24 hours. The sections are washed in probe wash buffer (Molecular Instruments) at 37°C for 10 minutes twice, 2×SSCT at room temperature for 10 minutes twice, then incubated in the probe amplification buffer (Molecular Instruments) at room temperature for 30 minutes. HCR fluorescent amplifier sets were first denatured in a PCR machine with a protocol of 90 seconds at 95°C and 25°C and then mixed in the probe amplification buffer to be a 1% amplifier solution. Sections were transferred to the 1% amplifier solution and incubated at room temperature for 24 hours. Lastly, sections were washed in 2×SSCT for 5 minutes 4 times, then mounted and imaged or continued with the immunohistochemistry procedure.

### Behavioral and electrical stimulation c-Fos experiments

Animals are kept in a sound-isolated chamber. Vocal activity was manually monitored for the first three hours in the morning (end of dark cycle) using SAP2011 recordings before sacrificing. Analysis of audio recordings was used to classify the animals into the no_vocal, call_only or song & call group.

For electrically induced c-Fos experiments, surgeries were performed at the end of the dark cycle to minimize background levels of c-Fos. Animals were anesthetized, and 30 μA 100 Hz current stimulation was used in RA or PAG for 2 seconds every 15 seconds for 30 minutes, after which animals were kept anesthetized for another 90 minutes before euthanasia.

### Statistical analysis

Data were analyzed through GraphPad Prism and python codes. Normality was tested using the Shapiro-Wilk Test. Data are presented as mean ± standard deviation. Statistical tests are described in the figure legends.

### Statistics and Reproducibility

Each experimental result was produced independently and/or combining at least 3 separate cohorts with similar results. Figures displaying viral expression, neural circuit tracing, FISH or immunohistochemical labeling are broadly representative of each experimental group.

---

**Extended Data Fig. 1.**
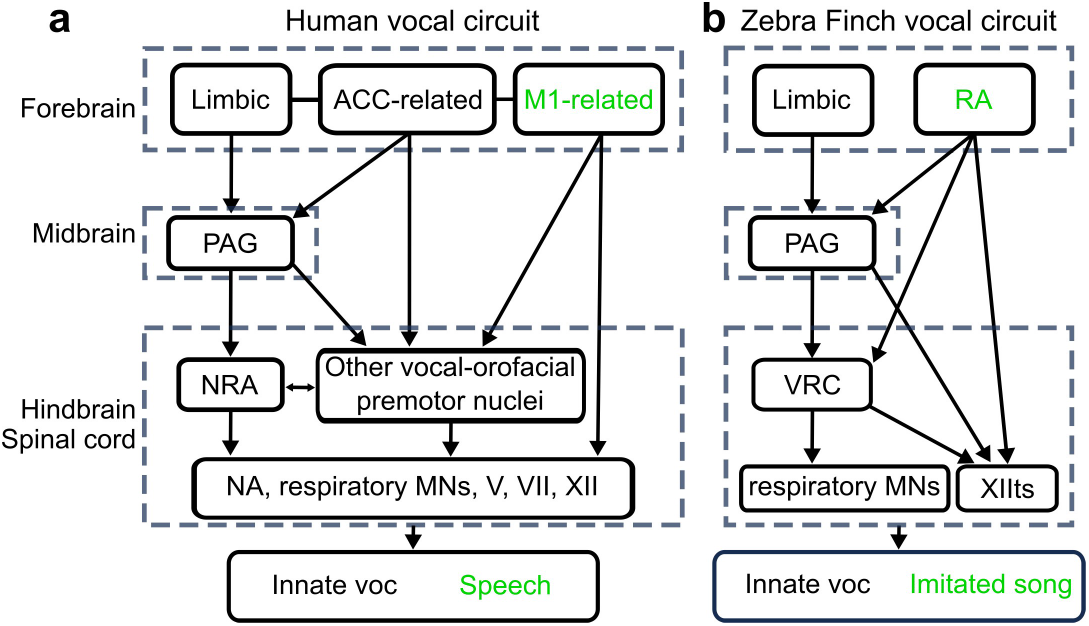
Descending vocal circuits. Detailed schematics of descending vocal circuits in human (a) and zebra finch (b).

**Extended Data Fig. 2.**
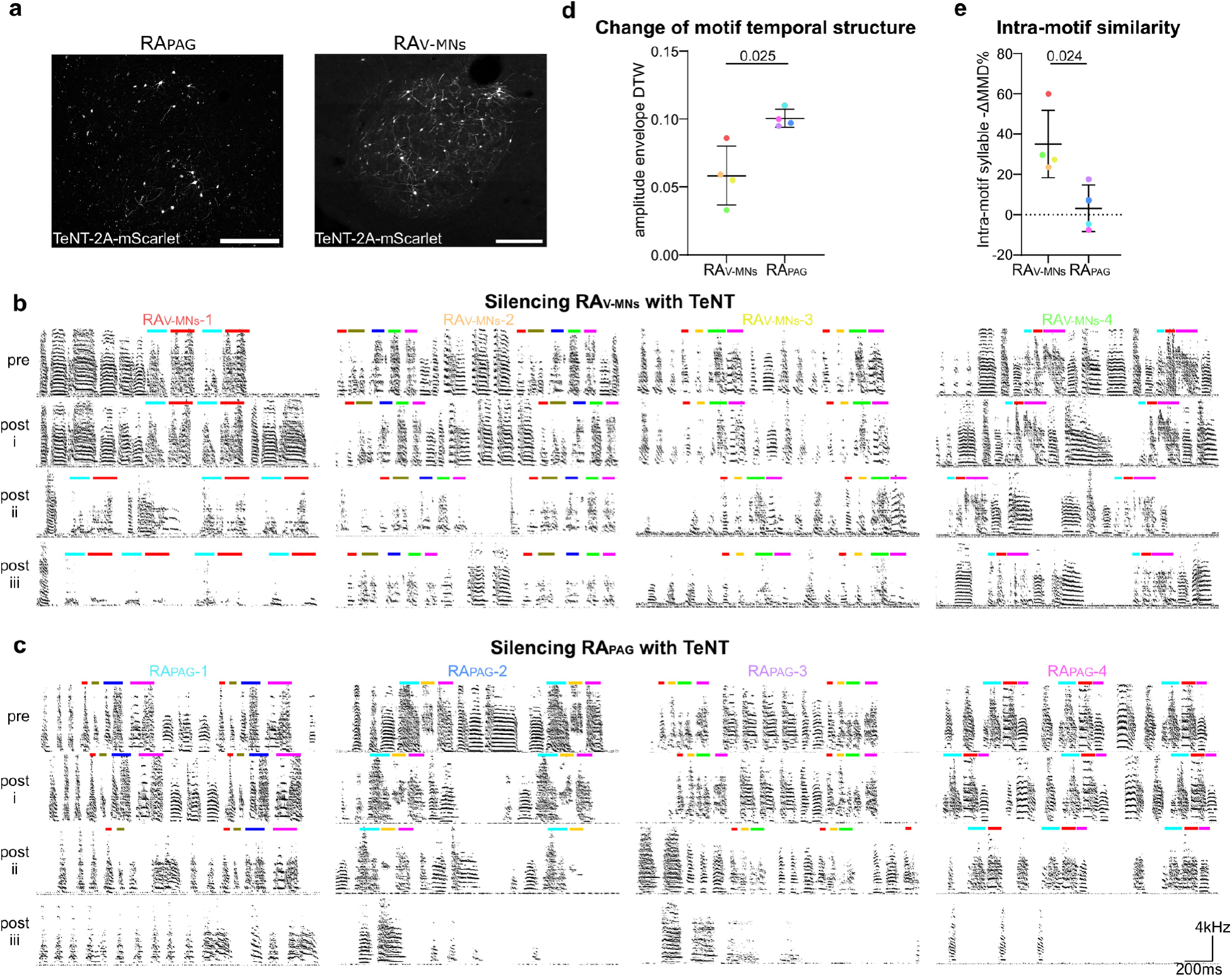
The role of descending pathways from RA in song production. **(a)** Retrograde viral expression in RA projection neurons. Left, PAG-projecting RA neurons. Right, V-MNs projecting RA neurons. Scale bar, 200 μm. **(b and c)** Progressive effect of silencing V-MNs/PAG-projecting RA neurons with TeNT. Example song spectrograms of 4 animals are shown from different timepoints of viral expression, pre and week 1,2,3 post viral injection. The animal names are color-coded to match the colors in panels d and e. Color bars mark syllables of each animal. **(d and e)** The same as Fig. 2c and d. Individual animals in RA_V-MNs_ and RA_PAG_ groups are labeled by different colors.

**Extended Data Fig. 3.**
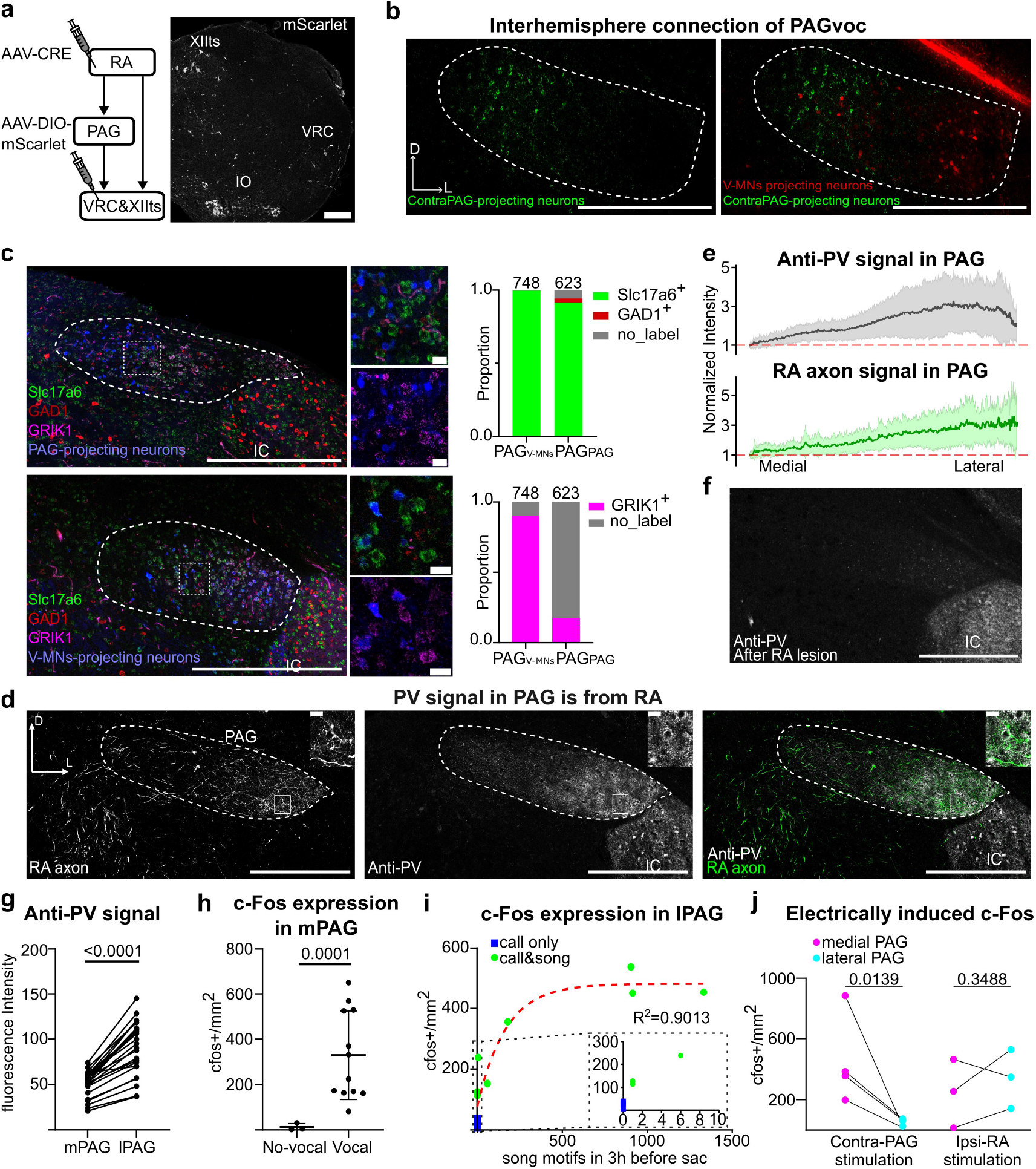
The vocal PAG is organized along the medial-lateral axis. **(a)** Anterograde viral jumping targeting hindbrain circuits. Left, schematic of viral injections. Right, expression patterns in the hindbrain. **(b)** PAG neurons projecting to the contralateral PAG are highlighted by the tracer in green. V-MNs projecting neurons are labeled in red. PAG is outlined by the dash line. **(c)** Cellular components of contraPAG-projecting neurons (top) and V-MNs-projecting neurons (bottom). Left, confocal images of FISH-Slc17a6, GAD1 and GRIK1 with fast blue labeling projections neurons. The dash box outlines the zoom-in image. Scale bar, 500 μm (20 μm for the zoom-in). Right, quantification of tracers-FISH signal colocalization. n=3 animals for each group. (**d)** Tracer-labeled RA axon and anti-PV signal in PAG, scale bar, 500 μm. PAG is outlined by the dash line. The solid square indicates position of the zoom-in on the top right, scale bar, 20 μm. **(e)** Distribution patterns of anti-PV and RA tracer signals in the medial-lateral axis of PAG. **(f)** PV signal in PAG after RA lesion, n=3 animals. Scale bar, 500 μm. **(g)** Medial and lateral PAG segmentation based on anti-PV signal. Mean fluorescence intensity of segmented lPAG and mPAG is quantified from brain slices of 4 animals. Welch’s t test. **(h)** c-Fos expression in the medial PAG after vocalization or without vocalization. n=3 for No_vocal group, n=12 for vocal group. Ratio paired t test. **(i)** Correlation of lPAG c-Fos expression and number of song motifs produced. Dashed lines outline the zoom-in window. An exponential plateau curve is fitted to the TeNT group data. **(j)** Quantification of electrical stimulation induced c-Fos expression in PAG. Ratio paired t test.

**Extended Data Fig. 4.**
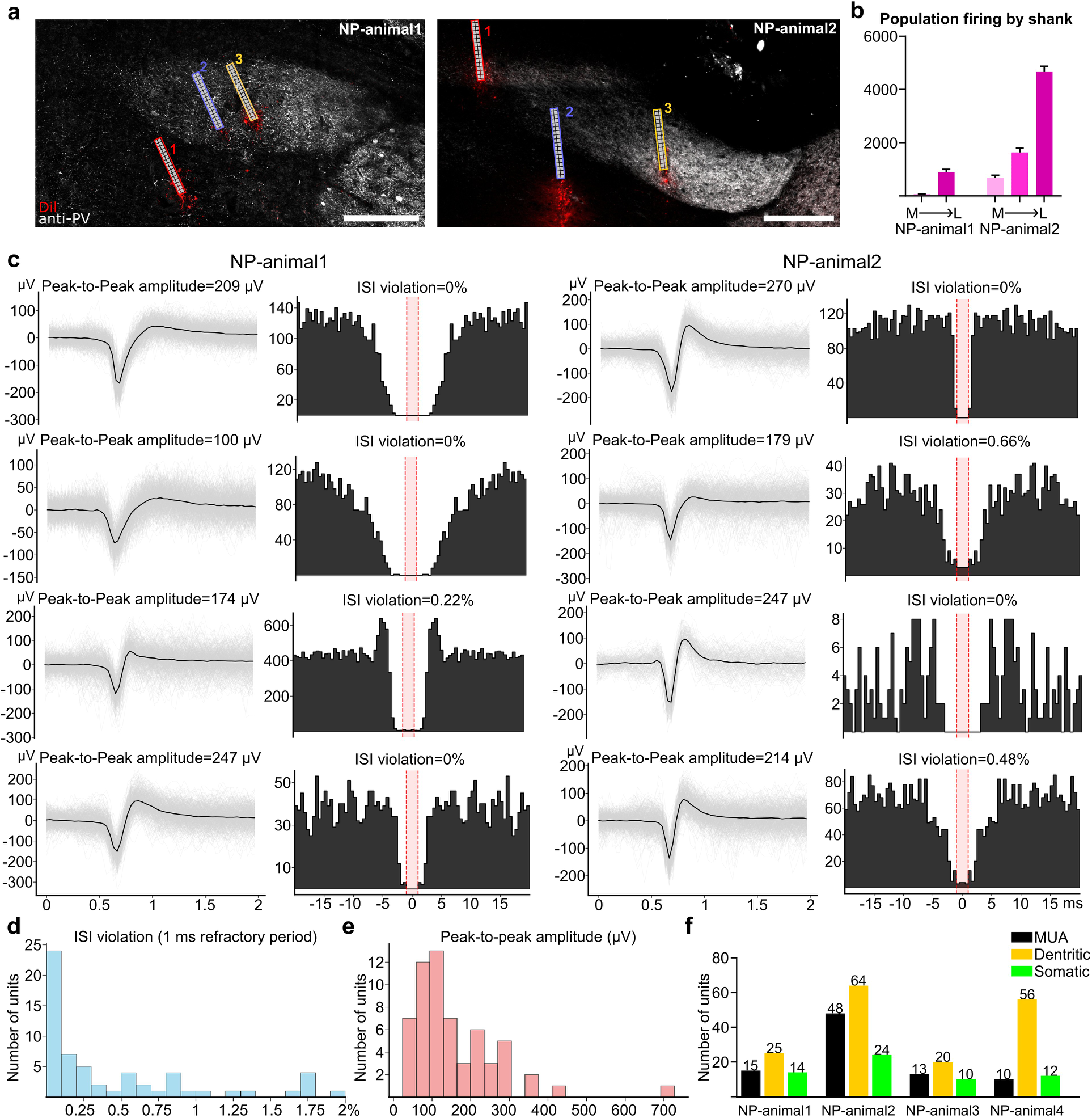
Neuropixels 2.0 recording in PAG. **(a)** Implantation tracks in PAG. PAG is visualized by anti-PV (grey), implantation tracks are marked by Dil (red). Scale bar, 200 μm. Left, recorded animal1; Right, recorded animal2. **(b)** Motif-modulated population firing rates grouped by each Neuropixels probe shank. 2 or 3 shanks are implanted into PAG in NP-animal1 or NP-animal2, respectively. Shanks that are inserted into more medial portions of PAG are labelled in light pink, while the lateral shanks are in dark pink. Variation between motif renditions is indicated by the S.D. error bar. **(c)** Example spike waveforms and autocorrelograms of 4 well-isolated somatic units from NP-animal1 and NP-animal2. In spike waveform plots, grey curves are individual spike waveforms, the black curves are the averaged waveforms. In autocorrelograms, red areas highlighted 1ms refractory periods, ratios of spikes violated the inter-spike interval (ISI) are calculated and showed on top of the autocorrelograms. **(d)** ISI violation histogram of well-isolated somatic units. **(e)** Peak-to-peak amplitude histogram of well-isolated somatic units. **(f)** Summary of all the vocal units from the four PAG recordings.

**Extended Data Fig. 5.**
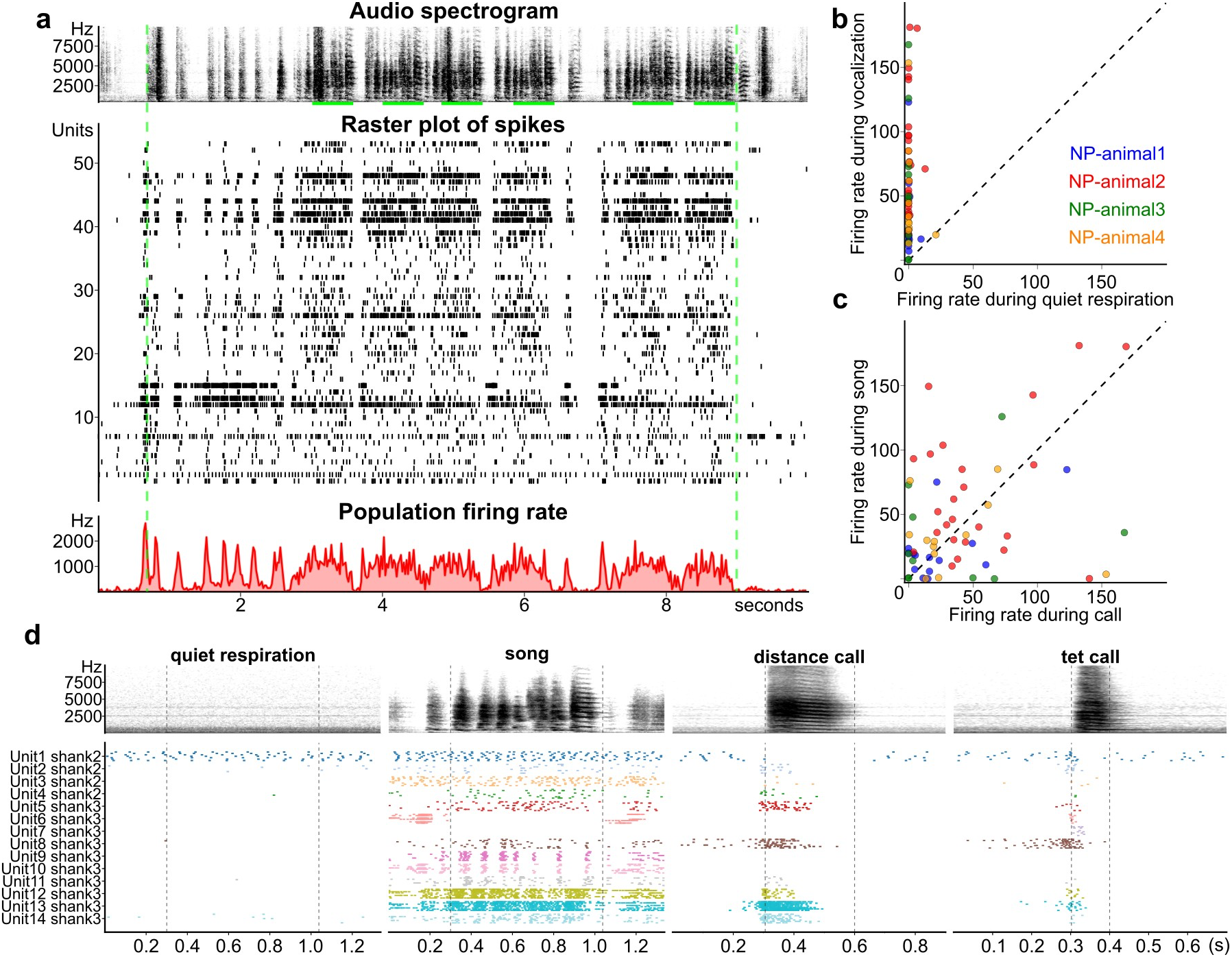
Neuropixels recording in PAG. **(a)** Synchronized song spectrogram, raster plot of sorted spikes and the population firing rate of NP-animal1. **(b and c)** Firing rates of well isolated somatic units during vocalization/quiet (b) and song motif/call (c). Units of different animals are labeled in colors. **(d)** Firing patterns of well isolated somatic units of NP-animal1 during different behaviors.

**Extended Data Fig. 6.**
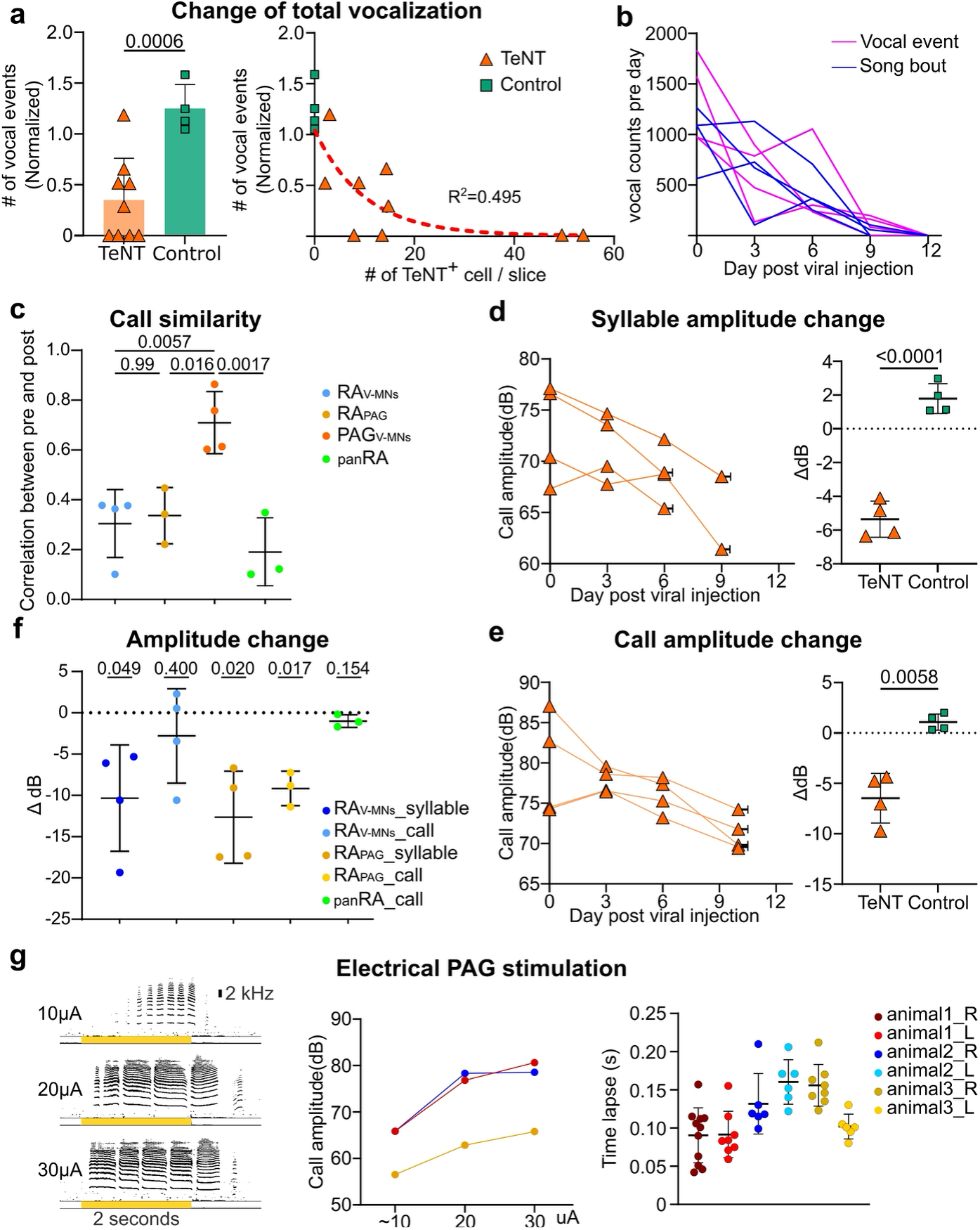
PAG_V-MNs_ are necessary for all vocalizations. **(a)** Change of total vocal event (including both song and call) after TeNT expression. **(b)** Number of song bouts and total vocal events pre and post TeNT viral expression of the four muted animals. **(c)** call similarity between pre and post TeNT in three TeNT groups and pharmacological panRA lesion group. One-way ANOVA and Tukey’s multiple comparisons test. **(d and e)** Change of syllable (d) and call (c) amplitude after TeNT expression in PAG compared with the control group. Welch’s t test. **(f)** Call and syllable amplitude change of RA_V-MNs_, RA_PAG_ and pan-RA lesion groups. n=3 for the RA_PAG_ group (one animal is excluded due to lack of distance call after viral expression), n=4 for the RA_V-MNs_ group, n=4 for the PAG_V-MNs_ group, n=3 for pharmacological panRA lesion group. One sample t-test with the theoretical mean at 0. **(g)** Naturalistic calls triggered by electrically stimulating PAG during anesthesia. Left, raw spectrogram of vocalization triggered by electrical stimulation. Yellow bars indicate 2 seconds electrical stimulation period. Middle, mean amplitude of calls triggered by different currents. Data from two hemispheres of the same animal (labeled by colors) are averaged based on stimulation intensity. Right, time-lapse between onset of electrical stimulation (30 μA) and first triggered calls.

**Extended Data Fig. 7.**
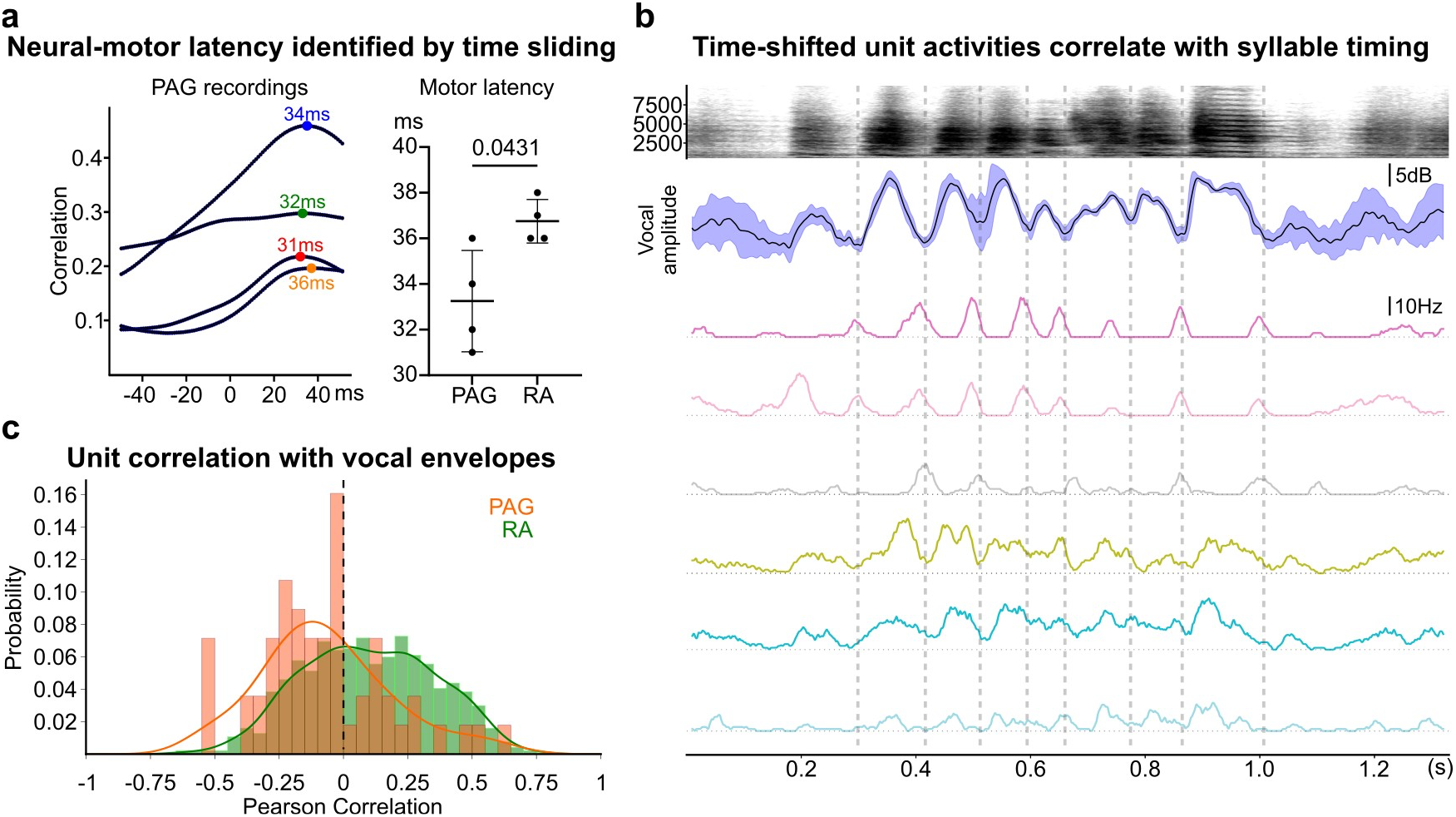
Neural-Vocal correlation in RA and PAG. **(a)** Neural-motor latency identified by time-sliding. Left, time sliding correlation curves of neural/behavioral correlation matrix of the 4 PAG recordings. The best fitting points are highlighted. Right, motor latency of RA and PAG recording. Welch’s t test. **(b)** Neural correlation with vocal respiration timing of example units from NP-animal1. Top, averaged spectrogram. Middle, the vocal amplitude curve. Bottom, unit firing curves, shifted by 34ms forward. The grey dashed lines highlighted putative inspiratory points in song motif. **(c)** Distribution of correlation coefficients between unit activity and vocal amplitude curves of RA and PAG.

## Notes

### Competing Interest Statement

The authors have declared no competing interest.

